# Persistence of microcystin production of *Planktothrix agardhii* exposed to different salinity concentrations

**DOI:** 10.1101/444000

**Authors:** Julia Vergalli, Audrey Combes, Evelyne Franquet, Stéphanie Fayolle, Katia Comte

## Abstract

Recent reports tend to predict the increase of harmful cyanobacteria in water systems worldwide due to the climatic and environmental changes, which would compromise water quality and public health. Among abiotic changes, the higher salinities are expected to promote the growth of some harmful species such as *Planktothrix agardhii*, which is known to build up blooms in brackish areas. Since *P. agardhii* is a common cyanotoxin producer (microcystin-producing), we investigated here the growth and tolerance of this species when exposed *in vitro* to a range of salinity levels, while assessing its microcystins variation and production in batch cultures during a time-frame experiment of 18 days. The study revealed a salt acclimation of the brackish *P. agardhii* that still produced microcystins in salty cultures while maintaining its growth ability in low to medium salinities (ranged from 0 to 7.5 g L^−1^). For higher salinity concentrations (10 to 12.5 g L^−1^), microcystins were still detected, while significantly lower growth rates were obtained during the exponential growth phase. This suggests that moderate to high salt ranges do not inhibit the microcystins production of *P. agardhii* at least for several weeks. Finally, the predicted remediation perspectives in a context of environment salinization assumed by environmental policies may be insufficient to eradicate this potential toxic cyanobacteria, especially when this species is already dominant in the waterbodies.

## INTRODUCTION

The massive occurrence and proliferation of cyanobacteria worldwide are serious issues as their bloom-forming abilities impair water quality (Twoney *et al.* 2002) in many ways (*i.e.* increasing turbidity, reducing biodiversity, leading to anoxia of the water column) and because some of common species are able to produce various toxic metabolites such as hepatotoxins and/or neurotoxins (Chorus and Bartram 1999). The most frequently found in waterbodies, including brackish areas (Sivonen and Jones 1999) are hepatotoxic microcystins (MCs) that can affect all living organisms from ciliates to fish (Combes *et al.* 2013; Ressom *et al.* 1994) and threaten human health (Chorus and Bartram 1999; Pouria *et al.* 1998).

MCs are cyclic heptapeptides that strongly (and irreversibly) inhibit serine-threonine protein phosphatases type 1 and 2A (Pearson *et al.* 2010) leading to cell disruption and death (Djediat *et al.* 2011). MCs have many structural variations (*i.e.* depending on the L-amino acid at the position 2 and 4 respectively from the whole MC architecture), and to date, over 200 MCs variants have been identified (Spoof and Catherine 2017) with different cytotoxic potentials; depending on the tested MCs variants (Shimizu *et al.* 2014). While reports on the biosynthesis and chemical processes of the MCs are constantly in progress, the forces underlying toxin production, *i.e.* the ecological and biological functions of MCs for the producing-cells still remain elusive and mostly contradictory (Babica *et al.* 2006). Various hypotheses for the possible role of MCs have been proposed, including: allelopathic effects (Leao *et al.* 2009) grazer defenses, light harvesting adaptation (Kaebernick and Neilan 2001). Recent findings suggest a possible involvement in intracellular processes and in primary metabolism (Zilliges *et al.* 2011), while excluding an essential role for growth (Hesse and Kohl 2001). Moreover, one of the most challenging questions is how environment influences the changes in MC concentration during cyanobacterial blooms. Indeed, a better understanding of the environmental factors triggering and/or involving the variations of the MC production and changes in the composition of toxic *vs* non-toxic cells, is highly required to help to predict the potential health hazards.

Numerous studies have shown that some environmental parameters may influence the MC production in toxic cells, including (i) the prevalence of toxic clones *vs* non toxic ones (Briand *et al.* 2005) during unfavourable conditions (Kurmayer *et al.* 2004), (ii) the increase of MC amount in toxic cells (Sivonen and Jones 1999) and (iii) changes in the MC variants composition (Tonk *et al.* 2005; Pearson *et al.* 2010). Among the possible causal factors are: the nutrient concentration (Downing *et al.* 2005), temperature and light (Wiedner *et al.* 2003), the iron concentration level (Sevilla *et al.* 2008) and pH (Song *et al.* 1998). Much less is known on many other abiotic parameters such as hydrologic variability, water bioavailability and salinity oscillations (N’Dong *et al.* 2014). Besides, the results are largely inconsistent as many factors (*i.e.* abiotic and biotic) may act in synergy and affect at different levels, the physiological state of the producing-cells (Davis *et al.* 2009).

All recent reports tend to predict that climatic change will exacerbate the dominance of harmful cyanobacteria in aquatic ecosystems worldwide (Paerl and Paul 2012; Carey *et al.* 2012). Indeed, the eutrophication heightened by human activities, coupled with environmental changes (such as rising temperatures, enhanced stratification of the water column) should trigger and increase the frequency, the biomass and the duration of the harmful cyanobacterial proliferations of specific species in waterbodies (Paerl and Otten 2013; Hagemann 2011; Fastner *et al.* 1999). With regards to global warming change, the oscillations in precipitation including episodic periods of intensive rainfalls (*i.e.* floods) *vs* droughts, could be effective events in expanding the bloom-forming species distribution along the freshwaters to the coastal areas (Lehman *et al.* 2005), especially if they are able to tolerate some moderate to high salt ranges. Thus, the rapid runoff including toxic cyanobacterial transport, may contaminate and thus impair the aquaculture and fisheries plants located in downstream waters (Robson et al. 2003; Preece et al. 2017).

*Planktothrix agardhii* (Gomont) Anagnostidis & Komàrek is one of the most common freshwater MCs producer in temperate areas (Chomerat *et al.* 2007) and has also been reported to produce some blooms in several brackish waters (*i.e.* from 3 to 11 g L^−1^ of NaCl) (Rojo *et al.* 1994; Villena *et al.* 2003). However, there are very few data on the influence of salinity on the MC production, because these widespread MC-producing species are mainly encountered in freshwaters. While *P. agardhii* is known to persist in brackish areas, it is important to investigate whether a rise of salinity may affect or not the ability of *P. agardhii* to produce MCs, as the actual remediation policy is performed by increasing the salinity of damaged and polluted waters to eradicate harmful organisms (Moisander *et al.* 2002; Von Alvensleben *et al.* 2013) (*i.e.* including potential toxic cyanobacteria) but for which evidence-based reports are still lacking. Therefore, we investigated the response of a dominant cyanobacterium *P. agardhii* strain originating from an oligohaline pond, to a gradient of salinity, and aimed to determine: i) the influence of the salinity range on the *P. agardhii* bloom development (growth and morphological changes), in batch cultures during a period of 18-20 days; ii) the influence of salinity on the effective MC production.

## MATERIALS AND METHODS

### Experimental section

#### Strain isolation and culture conditions

The *P. agardhii* strain (‘Brack’ strain) used in this study, was collected from the Olivier pond, in the vicinity of Istres, near the city of Marseille (in the south of France), located at 43° 30′ 46″ N latitude and 4° 59′ 17″ E longitude. The Olivier pond is a eutrophic and oligohaline waterbody (average salinity of 3 g L^−1^), covering an area of approximately 225 ha, with a maximum depth of 10 m. *P. agardhii* is the dominant cyanobacterium throughout the year (Vergalli 2013). Water samples were collected at the pond surface during a *P. agardhii* bloom in order to isolate filaments. After isolating a single filament (Rippka 1988), the strain was maintained for several years, under non-axenic conditions in Z8 liquid medium (Kotai 1972), at 22°C, using a light:dark cycle of 14:10h and a constant bubbling air to ensure homogeneous mixing and to provide sufficient quantities of inorganic carbon. The ‘Brack’ strain was assigned to the species *P. agardhii*, according to the morphological criteria provided by (Komárek and Anagnostidis 2005) and maintained in the Paris Museum Collection (PMC-MNHN) under the reference PMC1014.18

#### Experimental setup

For the experiments, NaCl was added to reach the final salinity concentrations of: 3, 5, 7.5, 10, 12 and 15 g L^−1^ and transferred with Z8 medium, into 250 mL Erlenmeyer flasks. The control corresponded to the culture maintained in Z8 medium (NaCl = 0 g L-1). Five replicates of each salinity concentration were checked with a conductivity meter (WTW LF330 Weilheim, Germany). The flasks were then inoculated with ‘Brack’ pre-culture in exponential growth phase and adjusted to obtain an initial OD_750_ = 0.1. Batch cultures were maintained in growth chambers under the same experimental conditions as described above. The flasks were regularly replaced in order to homogenize the light exposition provided in the growth chambers.

#### Growth measurement

##### Biomass and growth rate

The growth kinetics of ‘Brack’ strain cultures were monitored by measuring optical density at 750 nm using a Shimadzu UV-1700 spectrophotometer. The growth rate (μ) was calculated using the following equation:

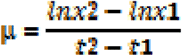

where t1 and t2 correspond to the measurement times (*i.e. t1*: the beginning of the exponential phase and *t2* (Day): the end of the experiment) x2 and x1 correspond to the biomass (expressed in OD values) at time t (with t2>t1).

##### Biovolumes and filaments’ length

The biovolumes (μm3) were assessed on the basis of the cylinder shape of filaments, according to (Sun and Liu 2003):
BV = 0,5Π lw

where l is the filament length (μm) and w the filament width (μm). A mean of 20 filaments was randomly measured in a counting chamber, using a micro-scale with a Nikon Labphoto2 microscope.

Fluorescence microscopy was performed with a Zeiss Primo Star microscope equipped with an AxioCam IcC1 cam. Epifluorescence images were recorded with specific filter (CY3) presets for chlorophyll *a* and acquired with the same time exposure set (AxioVision LE software).

#### Characterization of MC-variants

##### Template preparation

Three ml of mature culture of the ‘Brack’ strain were centrifuged (4000g for 10 minutes). The supernatant was discarded and the cell pellet was resuspended with 1 mL of methanol/water (90/10, v/v), followed by for 4 pulses of sonication on ice for 30 seconds. The mixture was centrifuged at 8000 g for 15 minutes at 4°C. The supernatant was collected, filtered (GF/C 1.2 μm) and evaporated (with a speed-vac concentrator at 40°C). The extract was dissolved in 100 μL water with 0.1% formic acid and centrifuged (4000 g × 5 minutes, 4°C). The supernatant was directly injected into the LC/ESI-MS system.

##### LC/MS analysis

LC/ESI MS and LC/ESI-MS/MS experiments were performed on a liquid chromatograph (LC) (UltiMate 3000®, Dionex) coupled to a Quadrupole-Time of flight (Q-TOF) hybrid mass spectrometer (Pulsar, Applied Biosystems) equipped with an electrospray ionization source (ESI). The chromatographic separation was conducted on a ACE3-C18 reverse-phase column (100 mm ×1 mm ×3 μm). Mobile phases were MilliQ water containing 0.1% (v/v) formic acid (A), acetonitrile containing 0.07% of formic acid (v/v) (B). The LC separation was achieved at a flow rate of 40 μl.min^−1^ using a gradient elution from 10 to 30% of solvent B in 5 minutes, then, from 30 to 70% B in 17 minutes, hold at 70% B for 5 minutes, return from 70 to 10% B in 3 minutes and hold at 10% B for 15 minutes. The mass spectrometer was operated with an electrospray ionization source in positive ion mode. For mass spectra, the capillary voltage was set to 2500 V with a declustering potential of 20 V. Full scan mass spectra were performed from 100 to 1500 m/z at 1s/scan in continuum mode. Fragmentation spectra were obtained in automatic mode using nitrogen as a collision gas, with collision energy automatically determined by the software according to the mass-to-charge ratio (m/z) values.

##### LC/ESI-MS et LC/ESI MS/MS data analyses

MS/MS spectra were analysed manually for highlighted spectra which contained the fragment ion (m/z= 135.1) characteristic of MCs fragmentation. All others ions fragments present on the fragmentation spectra were used to elucidate the structure of the MCs. The LC/ESI-MS data were processed using BioAnalyst 1.1 software. The molecular weight distribution of species (ranging from 100 Da to 1500 Da) observed in each sample were generated using the LC-MS reconstruct option. As the signal observed for both MC standards were close, the proportion of each variant was calculated by comparing the peak area corresponding to a given MC-variant to the total peak area of all MCs variants in a given sample.

#### MC concentration

Microcystins concentrations were determined by Enzyme Linked Immuno-Sorbent Assay (ELISA) using the MC-ADDA ELISA kit, (Abraxis LLC). ELISA tests were applied on supernatants from the cultures (*i.e.* cells pellets and supernatants), previously disrupted by a sonication on ice (2 pulses of 1 min, max. speed) according to the previous protocol Comes et al. 2013). The mixture was then centrifuged at 8000 ×g for 15 min at 4°C. The supernatant was collected and diluted in water v:v= 1:100 to 1:1000 (according to the biomass between T0 and T18) to avoid some matrix effects and potential salt interferences, as mentioned in the “Technical bulletin for microcystins in brackish and seawaters samples”(Abraxis). The measurements were performed in duplicate, on different samples exposed to each salinity treatment at Days 0, 2, 6, 8, 10 and 18. The limit of detection was approximately 0.10 ppb (μg L^−1^). The MC contents were expressed in μg L^−1^equivalent of MC-LR. Due to the positive correlation between the biovolumes (*i.e.* quantitative unit) and the biomass (*i.e.* OD_750_ values), (r2= 0.80; n=40, Fig. S2) the MC contents were converted and normalized per biomass (OD_750_) as a proxy of MC quota in order to compare the different MC patterns overtime by minimizing the growth factor.

#### Statistical analyses

‘Brack’ growth curves were fitted with the best trend approximation from absorbance measurements overtime, following the equation (Kahm *et al.* 2010):

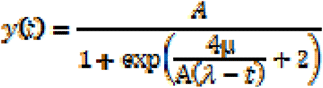

where ‘A’ is the asymptote in the curve and an estimation of the maximal density of the population reached during the life cycle;

‘μ’ is the maximal slope of the growth curve and characterizes the exponential growth phase (day 6 to 18); ‘λ’ is the lag-phase period of the growth (*i.e.* Day 2).

Two parameters (growth rate and maximal density) obtained from the logistic curves implemented with the ‘grofit’ package (Kahm *et al.* 2010) were used to compare the ‘Brack’ growth under various salinity treatments. Normality and homoscedasticity were systematically checked, using the Shapiro-Wilk and the Fligner-Killeen tests respectively. Consequently, the significant differences of growth (growth rate, biomass and filaments’ length) between the salinity treatments were performed by the One-Way Analysis of Variance (ANOVA) (n=5) and Tukey’s post-hoc test. The differences in MC concentrations were tested by Kruskal-Wallis test (between salinity treatments) and by Mann-Whitney test, when the MC patterns were compared to the control. The Pearson correlation coefficients were calculated between the growth variables and the MC concentrations. All statistical tests were carried out in R-2.14.0 environment and Statview (Roth et al. 1995).

## RESULTS

### Cell growth and morphological changes induced by exposition to various NaCl concentrations

Optical density (OD_750_) was recorded every day (from T0 to T18) to monitor the cultures growth and calculate the growth rate (μmax). While the highest salinity treatment (15g L^−1^) had a drastic effect on the strain growth (Fig. 1), the Brack strain was able to survive and grow from low to high salinity concentrations (from 3 to 12 g L^−1^) along the timeline of the experiment (18 days).

**Figure 1.**
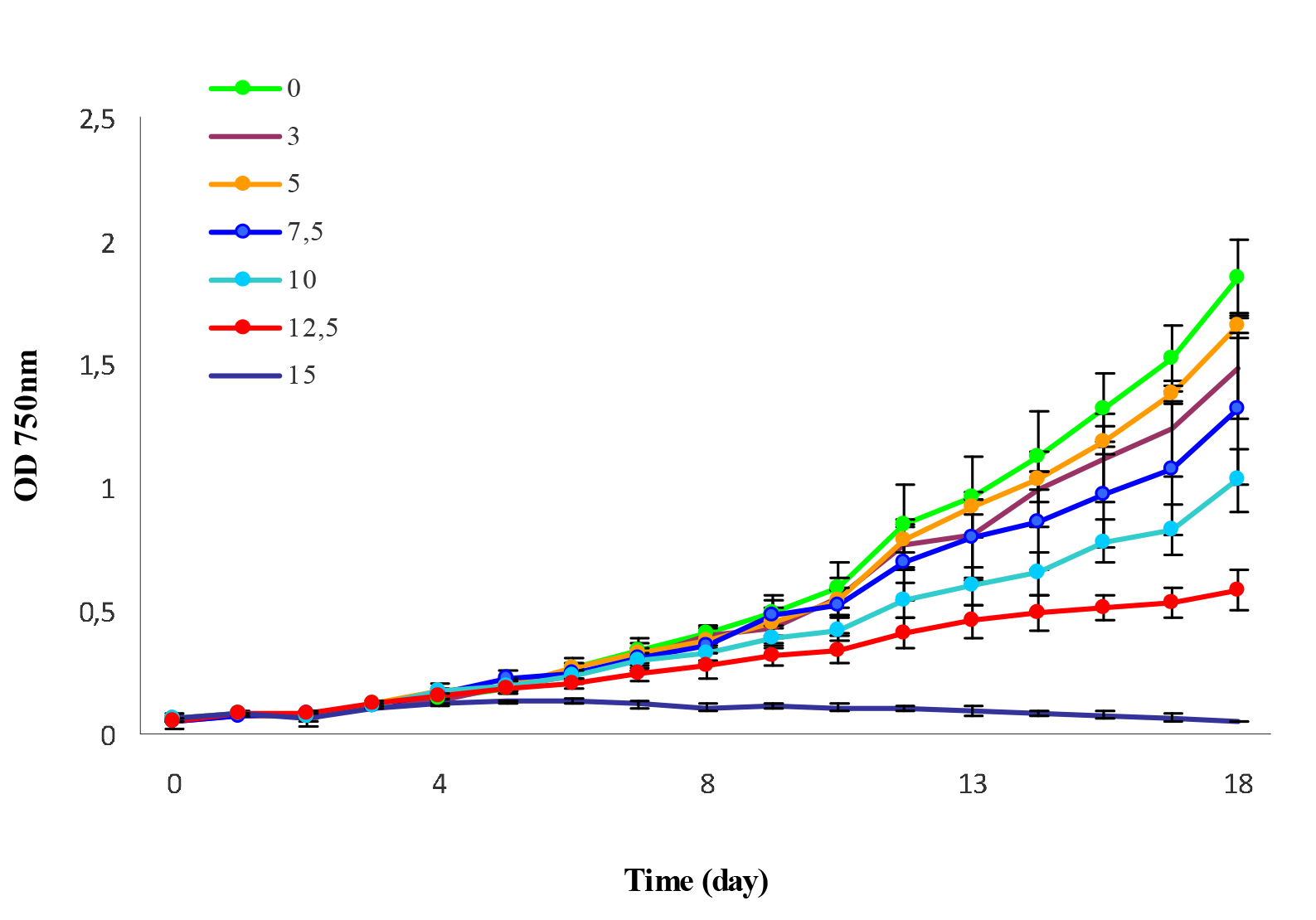
Growth dynamics of *P. agardhii* ‘Brack’ strain at different salinity concentrations, obtained using OD_750_ values (n= 5, ± SD) from T0 to T18 (A) and fitted with the ‘grofit’ package.

Two growth profiles can be distinguished: one that includes the control and the low salt concentrations that corresponds to a progressive increase of growth and a similar growth rate (ANOVA, P> 0.05); and a second profile corresponding to the high salt concentrations (7.5 to 12.5 g L^−1^) which revealed a significant decrease in terms of biomass, growth rate and density (Table 1) especially at Day 8.

**Table 1.**
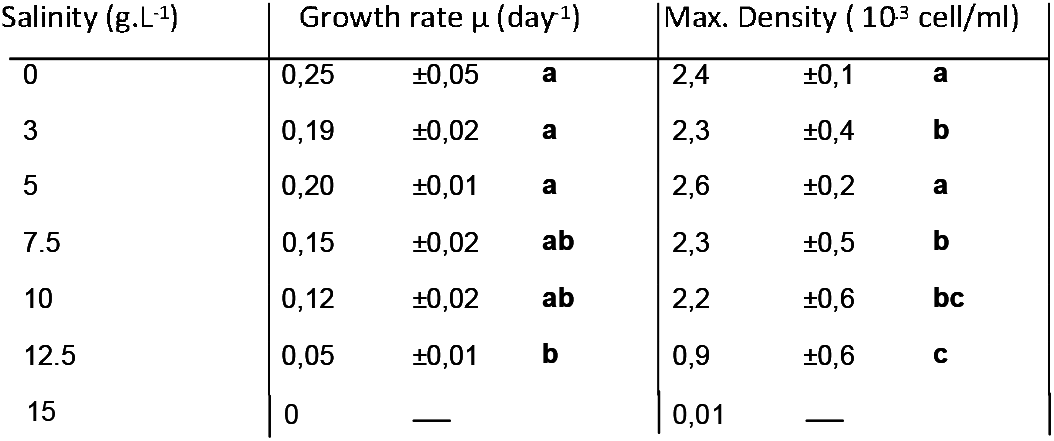
Descriptive parameters including the growth rate and cell density were determined from growth curves fitted by the ‘grofit’ model. The different letters mentioned above (a,b,c) indicate significant differences between the salinity treatments (Tukey test, p<0,01).

| Salinity (g. L<sup>-1</sup>) | Growth rate $\mu$ (day<sup>-1</sup>) | | | Max. Density ( 10<sup>3</sup> cell/ml) | | |
| --- | --- | --- | --- | --- | --- | --- |
| 0 | 0,25 | $\pm 0,05$ | **a** | 2,4 | $\pm 0,1$ | **a** |
| 3 | 0,19 | $\pm 0,02$ | **a** | 2,3 | $\pm 0,4$ | **b** |
| 5 | 0,20 | $\pm 0,01$ | **a** | 2,6 | $\pm 0,2$ | **a** |
| 7.5 | 0,15 | $\pm 0,02$ | **ab** | 2,3 | $\pm 0,5$ | **b** |
| 10 | 0,12 | $\pm 0,02$ | **ab** | 2,2 | $\pm 0,6$ | **bc** |
| 12.5 | 0,05 | $\pm 0,01$ | **b** | 0,9 | $\pm 0,6$ | **c** |
| 15 | 0 | — |  | 0,01 | — |  |

Additionally microscopic observations were performed every two days to estimate the size of filaments and detect some morphological changes in the whole cells and filaments. The physiological state of the filaments was assessed by the light and epifluorescence microscope taking intact morphology and chlorophyll autofluorescence intensity as indicators of survival. The decrease of chlorophyll content (*i.e.* OD values) with increasing salinity treatments, was correlated to a decrease of the total biovolumes *i.e.* consecutive to morphological changes and to a reduction of filament size (Fig. 2A). While the filaments had an approximate length of 240 μm at the beginning of the experiment (T=0), a first morphological variation was noted as early as Day 2 to Day 8 for moderate treatments (5 to 7.5 g L^−1^), consisting in a significant increase in the filament lengths up to a maximum (521μm as compared with the control (ANOVA p<0.05).

**Figure 2.**
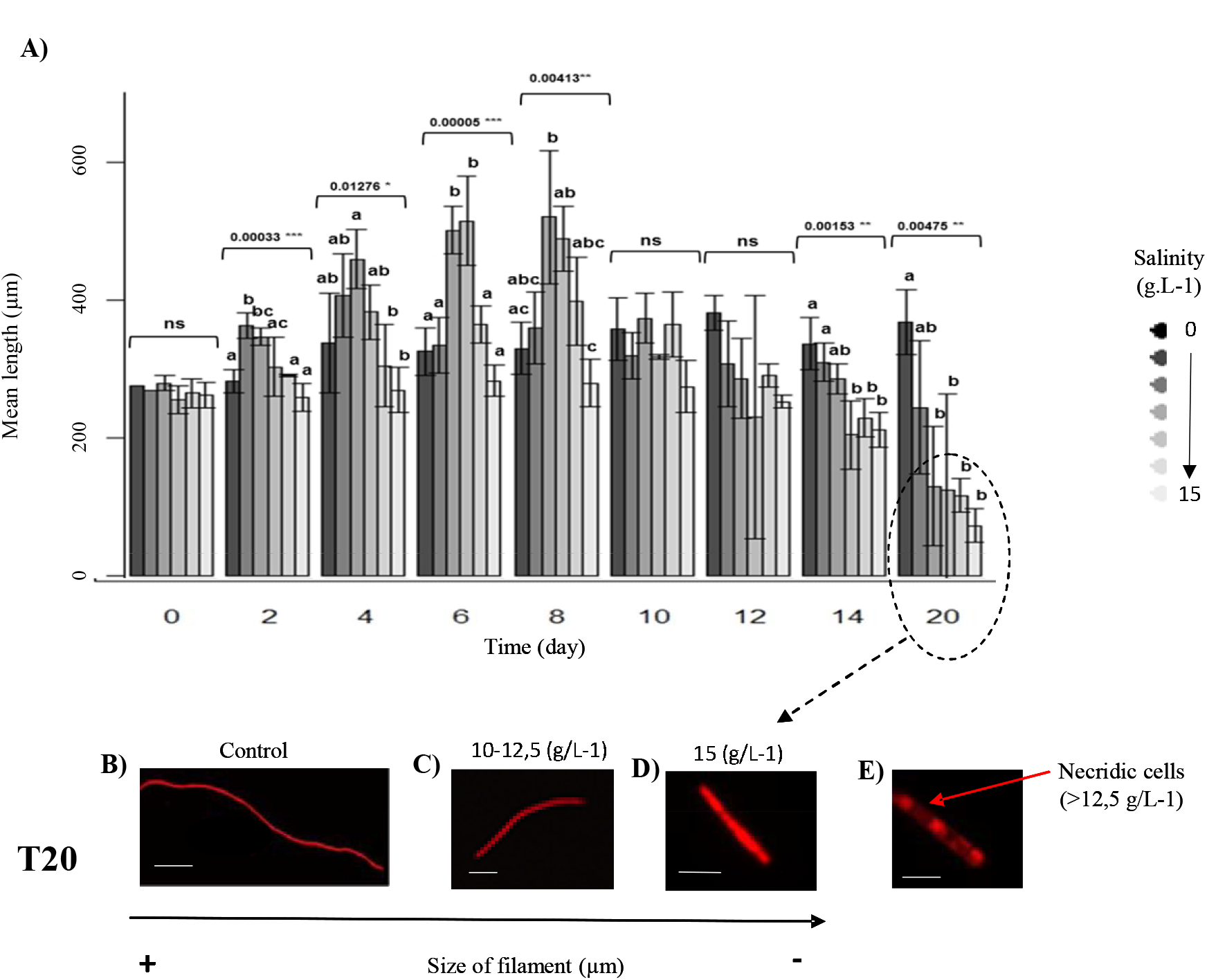
Variations of the filament length in μm, for cultures grown at different salt concentrations. A) Means with different letters (a, b, c) show significant differences between salinities concentrations (p < 0.05, ANOVA, Tukey post-hoc test). Error bars indicate standard deviation (n = 60). The asterisks indicate significant differences compared to the control (day =0) size. (NS= not significant difference; *= p<0.05; **= p<0.01; ***= p<0.001). B, C, D, E) Micrographs of size-type filaments observed in epifluorescence microscopy based on the chlorophyll autofluorescence (CY3 filter) in control (B), at 12,5 g L^−1^ (C) and 15 g L^−1^ of salinity (D, E) after 18 days of incubation. The yellow arrow (E) showed a necridia (non-fluorescent cell). Scale bars= 100μm (B); = 20μm (C, D) and = 10μm (E) respectively.

The elongation process was observed for cultures at 5 to 7.5 g L^−1^ of salinity, between Day 2 and Day 8 (ANOVA p<0,01), followed by a reduction of length beyond Day 10 (which were not significantly different from those measured for the control experiment-NS) (Fig. 2A). For the high salinity treatments (12.5 to 15g L^−1^), a significant reduction in the length of filaments was detected after 14 days, with a mean length not exceeding 60 μm as compared to the usual 350 μm of control experiment (Fig. 2 A). A remarkably high number of short fragments constituted by only 5-10 cells were observed at 15 g L^−1^ of salinity, at the end of experiment (Figs. 2D). Typical morphologies are shown in Figs. 2B, C, D. High intensity of the chlorophyll autofluorescence was still detected in short filaments, even after 20 days of incubation (Figs. 2C, D). Single cells located along the short filaments (at 12. 5 to 15 g L^−1^ of salinity) were sometimes completely dark (Fig. 2E- no fluorescent), corresponding to necridia or “suicidal-cell” referred to in [38] which split the filament in two fragments.

### Characterization of the MC-profile in the Brack strain

The characterization of the MC-variant composition was performed by liquid chromatography coupled to electrospray ionisation mass spectrometry (LC/ESI-MS/MS) in order to identify the chemo-type profile of the ‘Brack’ strain under optimal conditions (Fig. S1), as a various MC-diversity may exist within a same species Shimizu et al. 2014; Tonk et al. 2005). Five MC variants, (two major and three minor variants) were determined from cultured ‘Brack’ strain (Table 2).

**Table 2.**
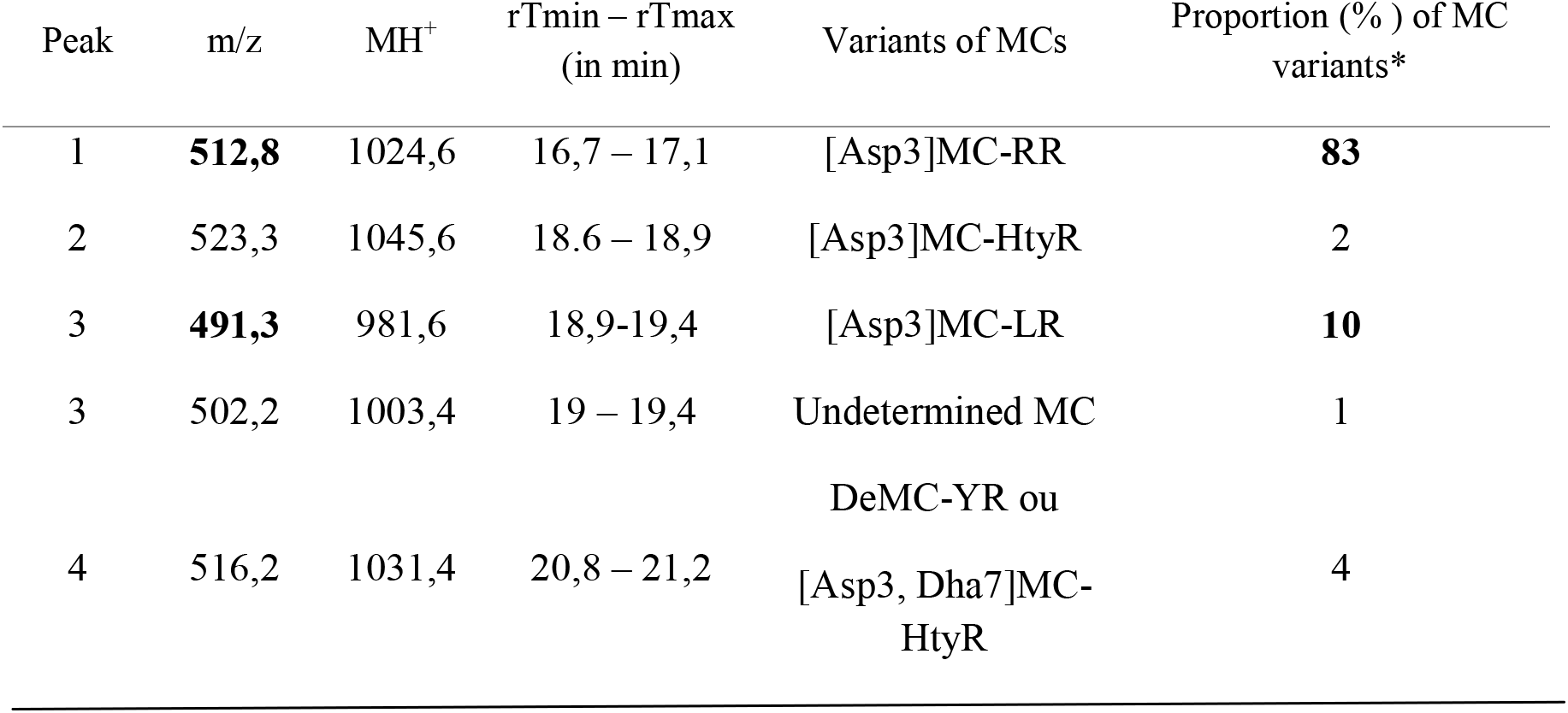
LC-ESI-MS determination of the individual MC-variants detected in the ‘Brack’ strain. The Identification of the MC-variant corresponds to ions detected on the mass spectra (m/z), and retention time (RT) compared to the standards. The proportion for each MC-variant was also included in the table. The m/z = 512.8; 491.3 and 523.3 with the respective retention time (RT) of 16.8 min; 19.2 min and 18.7 min were identified and confirmed by the corresponding MC standard. * Variants obtained under optimal conditions. Microcystins were quantified using [Asp3] MC-RR and [Asp3] MC-LR (Alexis Corporation) standards.

Based on the Adda fragment signal (m/z 135.1) and on the MS/MS spectra and the retention times of other ions (identical to MS standards and their MS/MS spectra), two major ions doubly charged [M+2H] ^2+,^ (Table 2) were identified respectively as [Asp3]MC-RR and [Asp3]MC-LR which altogether represent 93% of MCs present in this strain. Among the three minor ions, only one was clearly characterized as the demethylated [Asp3]MC-HtyR (Table 2), while the doubly charged [M+2H] ^2+^, (m/z = 516.2 with a RT of 21 min) was assigned to be either [Asp3]MC-YR or [Asp3,dha7]MC-HtyR. The ion m/z = 502.2 was undetermined and could not be assigned to [Asp3]MC-LR, referred to in (Yuan et al. 1999).

### Impact of the salinity treatments on the MC contents

In order to simplify the analysis and especially to get a better idea of the whole MCs present in the cell culture, the quantification of the total MCs, including both extra- and intracellular MC fractions, was assessed by using the microtiter plate MC-Adda ELISA test in all salinity treatments. The total MC contents (μg L^−1^) were positively correlated to the biomass (*i.e.* OD values) as suggested by the high r2 values for all the salinity treatments except 15 g L^−1^ (r2= 0,037), (Fig. 3). All the salts conditions, led to similar MC concentration profile including a progressive increase in MC content from 0 to Day 12, followed by a maximal concentration at Day 18 for 0 to 12,5 g L^−1^ of salinity (Kruskal Wallis, p>0.05). At 15 g L^−1^, a maximal peak was also present at Day 18, contrasting with a concomitant arrest in cell growth (μ=0).

**Figure 3.**
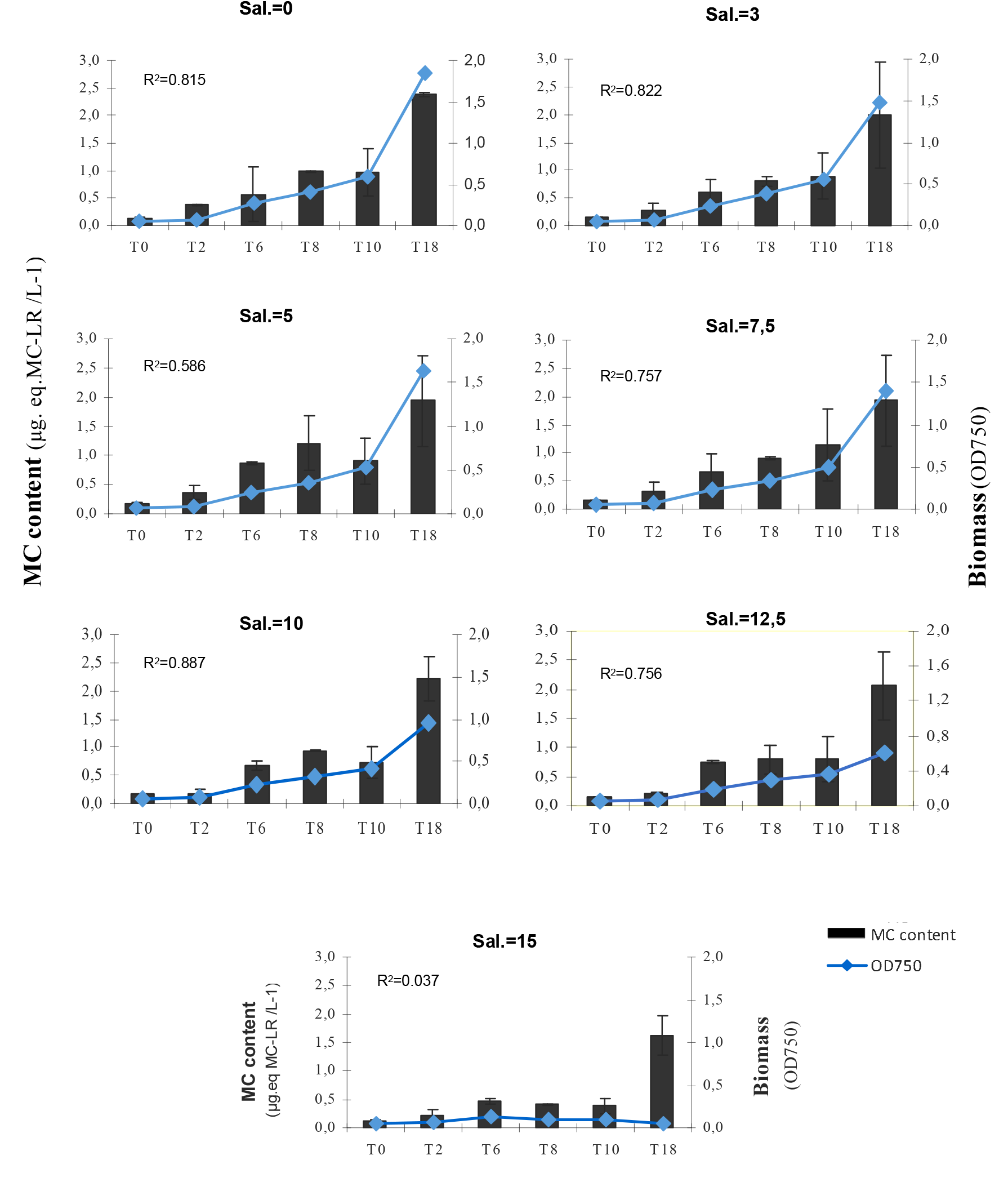
Variations of the MC contents (μg.eq. MC-LR/ L^−1^) of *P. agardhii* ‘Brack’ strain (means ± standard deviation, n=3) compared to the biomass (OD_750_ values) over time (18 days).

Since a positive correlation was obtained between the biovolumes (μm3) and the biomass (OD values- Fig. 2S) for all samples taken into consideration (r2= 0.85, n= 70), we could normalize the MC contents (μg equivalent) per biomass, as a proxy of MC quota to discriminate the MC profiles in various salinities and over time, so as to minimize the biomass factor. In more detail, the MC concentration showed four different profiles depending on the time frame and the salinity concentrations (two-way ANOVA, p<0.05). A first group was observed between 3 and 7.5 g L^−1^ and the control (Kruskal-Wallis, p>0.05) where the MC quota reached its highest value at Day 2 followed by a progressive decline from Day 8 to Day 18. Some slight differences were detected for the 5 g L^−1^ treatment (group 2) at Days 6 and 8 (p<0.05- Fig. 4), for which a still high MC value was noted at Day 2 but without the constant decline of MC previously observed in the group 1 and the control. For these groups, the MC quota was negatively correlated to the logarithm of biomass (r= −0.82, p= 5.8e-11, n= 42) and biovolumes (r= −0.76, p= 5.2e-09, n= 42) throughout the experiment (Days 2 to 18). A third MC profile was characterized for the 10 and 12,5 g L^−1^ treatments, including a rather stable MC concentration throughout the experiment with a moderate increase at Days 6 and 8 (Fig.4), that differed significantly from the control (p<0.01). The last group referred to the highest salinity (15 g L^−1^) and showed an increase of the MC concentration from Day 2 to Day 10, with a maximal value (7 times higher than control-p<0,001) at the end of experiment (Fig. 4).

**Figure 4.**
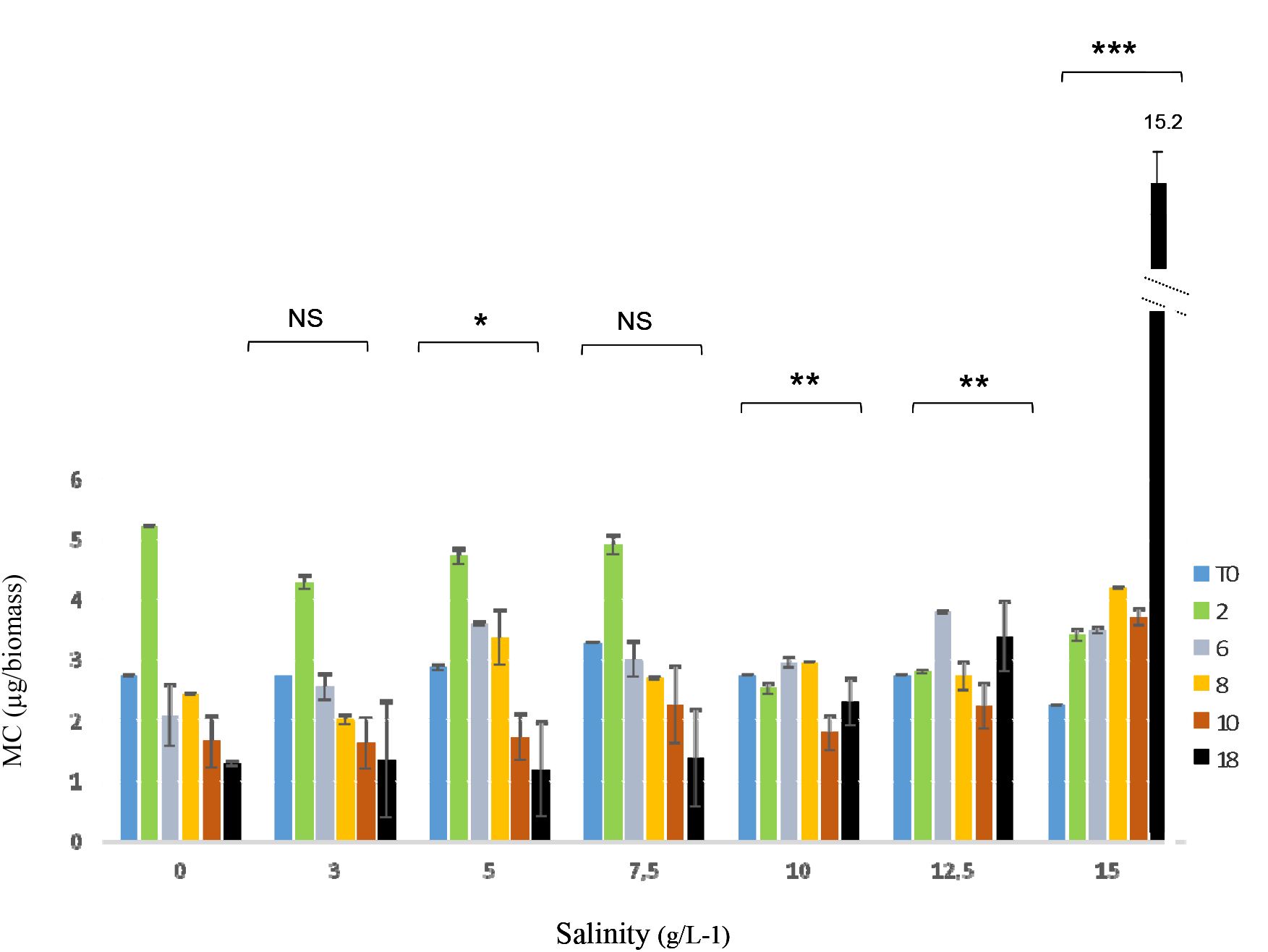
Evolution of MC contents (expressed as eq MC-LR μg/ biomass) in the ‘Brack’ strain, for each salinity treatment, overtime (0 to Day 18). The asterisks indicate significant differences compared to the control (Mann-Whitney test). NS= Not significant; *= p<0,05; **= p<0,01; ***=p<0,001).

As the ELISA test was applied to the whole culture (*i.e*. extracellular and intracellular fractions) it was not possible to confirm that the MC content came from the vivid cells or the media (suggesting a release of MCs after the cell death). Finally, the relative constant proportion of MC content from Day 2 to Day 10 for the 15 g L^−1^ treatment revealed no minute-lethal effect of the ‘Brack’ strain, suggesting a rather high tolerance of this strain to the salt stressor.

## DISCUSSION

### Effects of salinity on the growth and morphology of P. agardhii ‘Brack strain’

Our results have highlighted differences in the growth phases according to salt concentration, including a general decrease of the growth rate as the salinity increased (up to 7,5 g L^−1^), but at the same time, a persistence and a survival at 15g/L^−1 1^for several weeks. These findings reveal a higher tolerance of salinity for one *P. agardhii* strain when compared with the few available studies in literature (Chomerat *et al.* 2007; Komarek and Anagnostidis 2005; Orr *et al.* 2004). Nevertheless, morphological changes were observed throughout the experiment, suggesting an effective salt stress, which was still not sufficient enough to produce drastic effects on the survival of the population. While no significant difference was detected for the different salinities during the lag phase (p>0,05 between 0 to 2 days), a rapid increase in filament length occurred between 4 to 8 days for the cultures exposed to 10 and 12,5 g L^−1^ of salinity (Fig. 2). The elongation process could be the premise of cellular growth dysfunction or disruption in morphological processes (Singh and Montgomery 2013a,b), which cannot divide properly under this stressor. This temporary step was followed by a contrasting significant reduction of the filament length (Fig. 2A after 12 days) and the appearance of an increasing amount of short filaments (6-times less in size than control) from Day 10 to Day 18 (Fig. 2) for 10-15 g L^−1^ treatments. The presence of broken filaments suggests some cellular damages (Montgomery 2015) and has already been reported as a stress response of the cyanobacteria to environmental pressure (Singh and Montgomery 2013a; Poulickova *et al.* 2004). In these conditions, the shortened filaments could be a mechanism of defense used by the cells to preserve energy (Romo and Miracle 1993) and maintain the integrity of the few cells standing. Indeed, the *“in vivo”* chlorophyll autofluorescence revealed a high intensity of fluorescence signal in the short fragments at 15 g L^−1^ (Figs.2 C-D) which, combined with light microscopy, indicate that the integrity of the cells is still maintained as well as the active photosynthetic pigments within the cells. Although the best way to assess cell viability is the use of staining methods with fluorescent dyes (Pouneva 1997), autofluorescence of pigments may constitute a convenient method to evaluate the physiological state of the cell (Corrobé *et al.* 2017). At 12.5 to 15 g L^−1^ of salinity, intense fluorescence in most short fragments was recorded, while a few cells exhibit non photosynthetic fluorescent signals (dark cells- Fig. 2D) which correspond to necridia involved in the fragmentation of filaments (Komarek and Anagnostidis 2005; Castenholtz and Waterbury 1989). Interestingly, these suicidal cells can be seen as a relevant strategy for cell dissemination, consequently increasing the chance to find a more suitable habitat to restart a growing population (Komarek and Anagnostidis 2005).

### Effects of salinity on the total MC-content

In our investigation, the total MC content is positively correlated with the *P. agardhii* biomass for all salinity treatments (except the 15g L^−1^), which corroborates some previous studies (Paerl and Otten 2013; Mazur-Marzec *et al.* 2008; Dolman *et al.* 2012; Lyck 2004). No drastic effect was recorded immediately after exposure to high salinity, as expected for concentration (*e.g* 15 g L^−1^) which might release a massive MC amount within the first 24 hours due to the osmotic shock and the cell death, as was mentioned for other cyanobacterial species when exposed to pulse salt treatment (Tolar 2012). In this study, the MC-quota values were not significantly different between the 10-15 g L^−1^ suggesting that the cells are still able to cope with this stressor for a relative long period (from 2 to 15 days). Some studies have also shown that salinities up to 10 g L^−1^ do not affect the MC cell quota for *Microcystis, Anabaena* and *Anabaenopsis* genera (Black *et al.* 2011; Martin-Luna *et al.* 2015). Surprisingly, at the end of the experiment for the 15g/L treatment, a maximal peak of MC was detected (7- times higher than the control and 5 times higher than the previous MC amount at Day 15). Considering the cell density decline and the increased amount of short fragments, this suggests that massive cell disruption may have resulted in accumulation of stable MC in the medium. Indeed, some MC variants can be detectable and intact for up to several months (Zaspeta *et al.* 2014; Miller *et al.* 2010). The identification of the MC-profiles of the ‘Brack’ strain (performed by ESI-LC MS/MS) revealed that MC-LR is one of the two major variants, which is, with the dominance of Asp3 MC-RR, characteristic of the *Planktothrix agardhii* species (Fastner et al. 1999; Kurmayer *et al.* 2005). Because the ELISA tests were performed on both the extra- and intracellular MC fractions, it cannot be excluded that a possible high increase of MC production by living cells may contribute to the total amount of MC content.

### Ecological and management implications

Most of the investigations focusing on cyanobacterial responses to salt tolerance have reported controversial results (Tolar 2012; Tonk *et al.* 2007; De Pace *et al.* 2014) even at the intraspecific level (Otsuka *et al.* 1999). Some reports have shown discrepancies in the salinity thresholds for survival of *Microcystis* spp. (Orr *et al.* 2004; Tonk *et al.* 2007) for which some strains could resist to 10 g L^−1^ of NaCl, while other reached their limit at 2 g L^−1^, leading the authors to consider a strain-specific halotolerance rather than a species-specific trait (Orr *et al.* 2004). The salt-tolerance variability seems highly dependent on the life-history of each strain, implying direct and/or repetitive exposure to the stressor, which may induce acclimation and drive some intraspecific differences between strains. In our study, the unexpected survival of our *P. agardhii* strain to 15 g L^−1^ for several weeks may be a strain-specific response, as it was acclimated to the low salinity occurring in its brackish pond of origin (3 g L^−1^of salinity – Vergalli 2013). During the last decade, several episodes of increasing salinities (3 to 8 g L^−1^) were recorded in this pond, which may have selected some ecotypes adapted to the changing environment, as it suggested by Kirkwood et al. (2008). Common freshwater cyanobacterial species are able to tolerate salinity at low concentrations (Orr *et al.* 2004; Laamanen *et al.* 2001) and acclimate to salinity with time (Barron *et al.* 2002) and locations (brackish areas-Bergmann *et al.* 2008), unlike other phytoplankton groups (*i.e.* eukaryotes-Moisander *et al.* 2002). The potential shift of their halotolerance threshold and ability to tolerate salt variations arise the question of their potential persistence in the downstream waters including estuarine and coastal areas after meteorological-drifting events (caused by strong rainfalls or floods) (De Pace *et al.* 2014). It would be a serious issue since these cyanobacterial species are also toxin-producing cells and hence could contaminate the aquaculture and fisheries farms located along the freshwater-marine continuum (Bergmann et al. 2008). Finally, it may be a crucial issue for water management strategies based on the increase and/or oscillations of salinity concentration in freshwater systems, as already implemented in several countries such as the Netherlands (Verspagen *et al.* 2006). Our results clearly show that increasing the salt concentration of a brackish Mediterranean pond by water input or by a pseudo-natural salinization (Cf. Vergalli 2013) will not eradicate *Planktothrix agardhii* populations if the salinity is not up to 15 g L^−1^. Besides salt stress often increase lysis of MC-producing cells which may affect directly or indirectly all the living organisms in aquatic systems. Thus, care must be taken when considering increasing of salinity as a potential water management or remediation strategy. It may render a regular checking and security procedure necessary, especially in the recreational areas.

## CONCLUSION

Elevated salinities (up to 12.5 g L^−1^) affected the cellular growth and morphology of the Brack strain, as suggested by a lower growth rate and an increase of short broken filaments. However, *P. agardhii* was able to tolerate moderate to high amount of salinities. The threshold for normal growth seemed to be set at 15 g L^−1^ of salinity, but this concentration allowed survival of the strain, without a minute lethal salt-shock at least during the time frame (18 days). The constant amount of MC products overtime may lead to a real harmful effect on the environment and aquatic organisms. Our findings may be important to take into account when considering the water management policies based on salinity increase, planned by several countries to eradicate toxic and bloom-forming species. This study emphasizes the crucial need to further investigate the gradual and repetitive increasing of salinity as an indirect consequence of the global warming change, on the acclimation and/or adaptive response of these filamentous toxic cyanobacteria worldwide.

## ACKNOWLEDGEMENTS

Financial support was provided by the “Region Provence Alpes Côte d’Azur” (PACA-France). Many thanks to the mass spectrometry facilities of MNHN (Paris-France) and the technical coordinators from SIBOJAÏ.

## AUTHOR CONTRIBUTIONS

SF, KC, JV conceived and designed the experiments. JV and AC performed the experiments. SF and KC wrote the paper. All authors approved the final manuscript.

